# Integrated Proteomics Identifies Troponin I Isoform Switch as a Regulator of a Sarcomere-Metabolism Axis During Cardiac Regeneration

**DOI:** 10.1101/2023.10.20.563389

**Authors:** Timothy J. Aballo, Jiyoung Bae, Wyatt G. Paltzer, Emily A. Chapman, Rebecca J. Salamon, Morgan M. Mann, Ying Ge, Ahmed I. Mahmoud

## Abstract

Adult mammalian cardiomyocytes have limited proliferative potential, and after myocardial infarction (MI), injured cardiac tissue is replaced with fibrotic scar rather than with functioning myocardium. In contrast, the neonatal mouse heart possesses a regenerative capacity governed by cardiomyocyte proliferation; however, a metabolic switch from glycolysis to fatty acid oxidation during postnatal development results in loss of this regenerative capacity. Interestingly, a sarcomere isoform switch also takes place during postnatal development where slow skeletal troponin I (ssTnI) is replaced with cardiac troponin I (cTnI). In this study, we first employ integrated quantitative bottom-up and top-down proteomics to comprehensively define the proteomic and sarcomeric landscape during postnatal heart maturation. Utilizing a cardiomyocyte-specific ssTnI transgenic mouse model, we found that ssTnI overexpression increased cardiomyocyte proliferation and the cardiac regenerative capacity of the postnatal heart following MI compared to control mice by histological analysis. Our global proteomic analysis of ssTnI transgenic mice following MI reveals that ssTnI overexpression induces a significant shift in the cardiac proteomic landscape. This shift is characterized by an upregulation of key proteins involved in glycolytic metabolism. Collectively, our data suggest that the postnatal TnI isoform switch may play a role in the metabolic shift from glycolysis to fatty acid oxidation during postnatal maturation. This underscores the significance of a sarcomere-metabolism axis during cardiomyocyte proliferation and heart regeneration.

## INTRODUCTION

Heart failure is the leading cause of death in the U.S. and worldwide^1^. Despite its prevalence, there are no curative therapies that address heart failure’s underlying pathophysiology. After a cardiovascular ischemic insult that leads to sustained myocardial damage, such as a MI, the adult mammalian heart lacks the capacity to repair itself, leading to pathological cardiac remodeling and systolic heart failure^2^. Remarkably, the neonatal mouse heart possesses a striking capacity to regenerate after cardiac injury largely through the proliferation of pre-existing cardiomyocytes (CMs)^3,4^. During postnatal maturation, there is a switch in CM sarcomere isoforms as well as a metabolic transition from glycolysis to oxidative phosphorylation^5,6^. During these transitions, the mouse heart loses its capacity to regenerate as the sarcomeres undergo structural and functional maturation^6^, preventing sarcomere disassembly that is necessary for successful CM proliferation^7,8^. Additionally, the metabolic shifts in mature CMs lead to an increase in reactive oxygen species (ROS) production, which also contributes to CM cell cycle arrest^9,10^. However, the interplay between sarcomeres, the postnatal metabolic switch from glycolysis to oxidative phosphorylation, and cardiac regenerative potential remains poorly defined.

At the sarcomeric level, many different immature isoforms transition to their mature variants during fetal and postnatal cardiac development^5,11^. Specifically, Troponin I (TnI), the inhibitory subunit of the troponin complex, is crucial for Ca^2+^ modulation of CM contraction and relaxation^12^. Interestingly, TnI undergoes a transition from the immature ssTnI isoform to its mature cTnI isoform, which takes place during postnatal maturation and coincides with the loss of the regenerative capacity of the neonatal mouse heart^3,5^. Previous studies demonstrate that ssTnI increases Ca^2+^ binding affinity to troponin whereas cTnI results in lower Ca^2+^ sensitivity; thus, ssTnI overexpression leads to impaired CM relaxation kinetics under baseline conditions^13^. Remarkably, overexpression of ssTnI provides metabolic protection against ischemia. This is achieved primarily through enhanced glycolytic metabolism, likely driven by increased AMPK activity^14^. Furthermore, ssTnI overexpression improves the response to pressure overload injury, also due to elevated glycolytic metabolism^15^.

Recent evidence demonstrates that rewiring cardiac metabolism, whether by modifying mitochondrial substrate utilization, inhibition of succinate dehydrogenase, or inhibition of fatty acid oxidation, can promote adult CM proliferation and heart regeneration^16–18^. Although previous studies suggest a potential role for ssTnI in regulating cardiac metabolism, it remains unclear whether ssTnI regulates the postnatal metabolic switch and the cardiac regenerative capacity. Therefore, it is crucial to adopt an unbiased approach to understanding cardiac development and the interplay between sarcomeres and metabolism during regeneration.

High-resolution mass spectrometry (MS)-based proteomics provides an avenue to comprehensively and quantitatively define the alterations in protein expression that occur at the sarcomere and metabolic level throughout postnatal cardiac development, endogenous cardiac regeneration, and the cardiac injury response. Namely, there are two MS-based approaches commonly used to interrogate the proteome: the bottom-up approach in which proteins are digested into peptides prior to analysis and the top-down approach in which proteins are analyzed in their intact state. While bottom-up proteomics is commonly used for analyzing entire proteomes due to the ease of separating, ionizing, and fragmenting peptides, it faces challenges in peptide to protein inference and in mapping and quantifying post-translational modifications (PTMs). On the other hand, top-down intact protein analysis provides unambiguous proteoform identification and quantification of isoforms PTMs, but it is better suited for targeted sub-proteome characterization. In this study, we employ a combination of bottom-up and top-down proteomics, cardiac injury models, and histological analysis to comprehensively define the shifts in the proteomic landscape throughout postnatal mouse cardiac development and analyze the response to cardiac injury and regeneration. Specifically, we investigate how ssTnI overexpression influences baseline proteome composition and modifies the cardiac injury response through regulation of the metabolic protein signature.

## RESULTS

### Proteomic analysis of postnatal mouse heart development reveals alterations in developmental processes, metabolic proteins, and sarcomere composition

The neonatal mouse heart can fully regenerate following cardiac injury, but this regenerative capacity is lost by postnatal day (P) 7^3^. To understand the molecular basis of this regenerative capacity, we sought to determine how the cardiac proteome is remodeled during postnatal mouse heart development. To do so, we harvested mouse hearts at P1, P8, and P28 (n = 5 per group), and then analyzed the ventricular tissue using global bottom-up proteomics and targeted top-down proteomics of the cardiac sarcomere **(Figure 1A)**.

**Figure 1.**
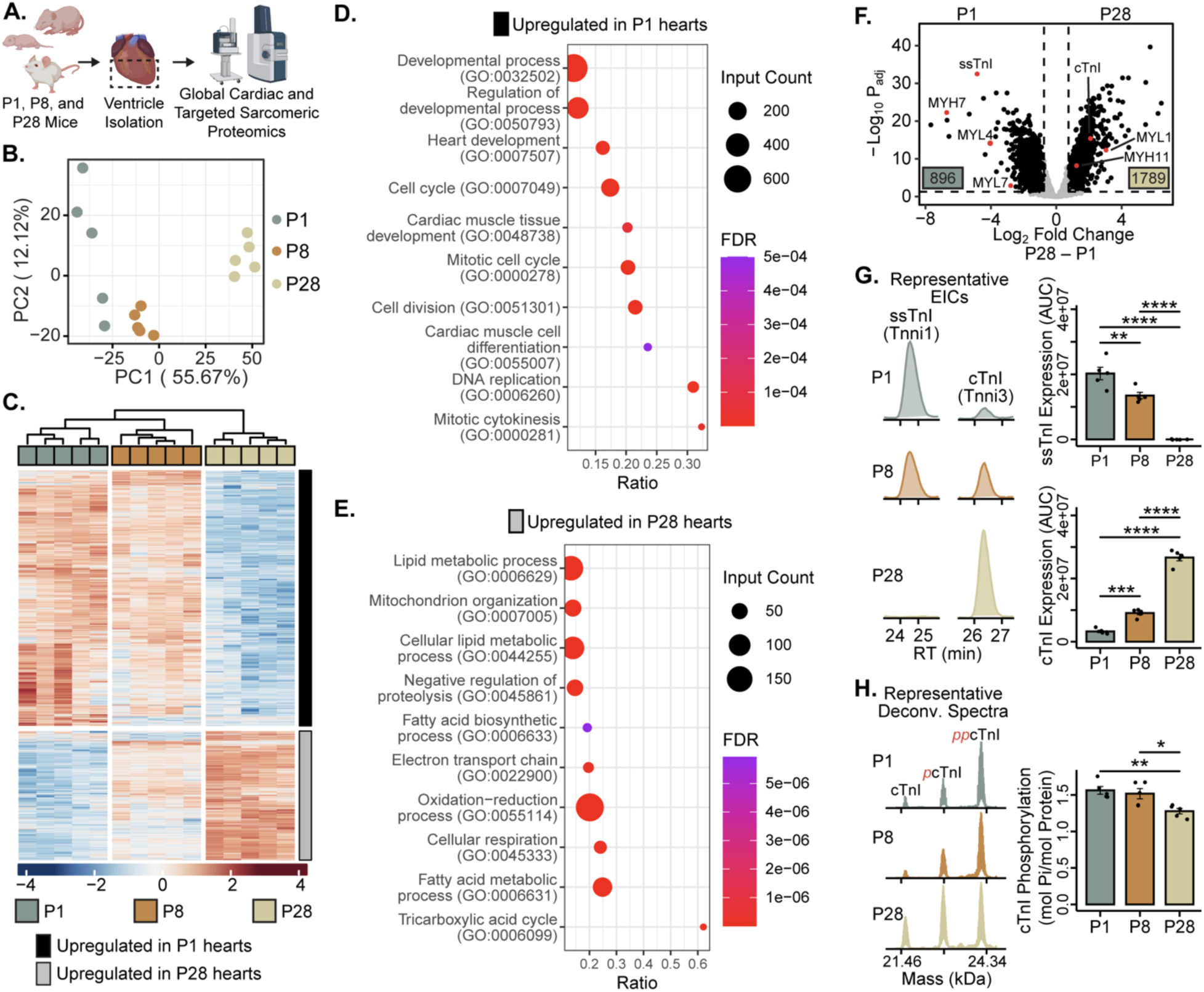
Proteomic analysis of postnatal mouse heart development reveals alterations in developmental processes, metabolic Proteins, and sarcomere composition. **A.** Workflow of experimental design. Mice were sacrificed at postnatal days (P) 1, 8, and 28 (n = 5 per group), the ventricles were isolated, and samples were prepared for global bottom-up cardiac proteomics and targeted top-down sarcomeric proteomics. **B.** Principal component analysis (PCA) of per-sample Log_2_ protein abundances demonstrates reproducibility among samples and separation among groups. **C.** Hierarchal heatmap (k-means columns set to 3, k-means rows set to 2) displaying z-score normalized protein intensities of all significantly differentially expressed proteins throughout postnatal development (adjusted p-value ≤ 0.05 and |Log2 Fold Change| ≥ 0.75 significance thresholds). The rows are separated into two clusters: proteins with high expression at P1 (black) and proteins with high expression at P28 (grey). **D and E.** Selected STRING biological process gene ontology (GO) plots of the proteins that are upregulated in P1 hearts (**D**) or upregulated in P28 hearts (**E**). Ratio represents the fractions of all proteins in the GO category that were identified. Dot size corresponds to the number of identified proteins within a GO category. Color represents FDR-adjusted p-value of the overrepresentation test. **F**. Volcano plot demonstrating fold-change in protein expression between P1 and P28 hearts. The number of significantly upregulated proteins per group is shown in the bottom corners of each comparison (n = 5 per group). Sarcomeric proteins that change throughout postnatal development are highlighted in the plot (adjusted p-value ≤ 0.05 and |Log2 Fold Change| ≥ 0.75 significance thresholds). **G**. Top-down proteomic analysis of ssTnI (Tnni1) and cTnI (Tnni3) expression throughout postnatal development. Left: Representative extracted ion chromatograms of ssTnI and cTnI. Right: Expression level analysis of ssTnI and cTnI. **H**. Top-down proteomic analysis of cTnI phosphorylation throughout postnatal development. Left: Representative deconvoluted spectra of cTnI. Right: Quantification of relative expression of phosphorylated cTnI to total cTnI expression. Bars represent average, error bars represent standard error of the mean (S.E.M.), dots represent individual data points. * denotes *p* ≤ 0.05, ** denotes *p* ≤ 0.01, *** denotes *p* ≤ 0.001, and **** denotes *p* ≤ 0.0001 as determined by one-way ANOVA followed by Tukey’s HSD.

In our global proteomic analysis of postnatal cardiac development, we quantified over 7,000 protein groups in all three biological groups **(Figure S1A, Table S1)**. Interestingly, fewer protein groups were quantified in the P28 hearts, which coincides with increased cardiac tissue maturation and was found to also occur during postnatal swine heart development^19^. Furthermore, among these 7,000 protein groups, there was a core proteome of 5,361 protein groups shared among all samples and a unique set of protein groups quantified at each timepoint **(Figure S1B)**. Additionally, the robust clustering in our principal component analysis (PCA) **(Figure 1B)**, where samples were separated by time along PC1 (55.67% of the variation), and the low variation in protein group intensity among biological replicates **(Figure S1C)** lent high confidence to downstream statistical analyses.

Using a limma-based differential expression analysis, we were able to identify over 2,500 differentially expressed proteins (DEPs) among P1, P8, and P28 hearts **(Table S2)**. Using these DEPs, we created a hierarchal heatmap and identified two strong trends **(Figure 1C)**: a group of proteins upregulated in P1 hearts, and a group of proteins upregulated in P28 hearts. Interestingly, P8 hearts displayed an intermediary protein expression profile, indicating a gradual transition in protein expression throughout postnatal development. Gene ontology (GO) analysis of the proteins identified in these clusters revealed biological processes related to development (GO:0050793, GO:0007507, GO:0055007) and mitosis (GO:0007049, GO:0051301, GO:0000281) upregulated in P1 hearts **(Figure 1D)**, highlighting the proliferative capacity of P1 hearts that supports endogenous cardiac regeneration. Conversely, GO analyses of the proteins upregulated in P28 hearts revealed processes related to lipid metabolism (GO:0006629), oxidation-reduction (GO:0055114), and negative regulation of proteolysis (GO:0044255) **(Figure 1E)**. Interestingly, the heart’s metabolic state plays a key role in its ability to regenerate after injury, with glycolytic metabolism promoting proliferation and fatty acid oxidation leading to cell cycle arrest^9,20^. This global analysis highlights how the proteomic landscape changes from a more proliferative to a terminally differentiated state during postnatal development. During this metabolic transition, there was also a distinct shift in sarcomere composition **(Figure 1F)**, with many immature sarcomeric proteins (TNNI1 (ssTnI), MYH7, MYL4, and MYL7) transitioning to their mature isoforms (TNNI3 (cTnI), MYL1, and MYH11). Interestingly, the ssTnI to cTnI isoform transition was among the most significantly differentially expressed proteins identified between P1 and P28 hearts **(Figure 1F and S1D)**. Using a highly quantitative targeted top-down proteomics analysis of the cardiac sarcomere **(Figure S1E-G)**, we were able to robustly quantify this change in troponin I (TnI) expression and found that by P8, there was about equal expression of ssTnI and cTnI, and by P28, ssTnI was completely replaced by cTnI **(Figure 1G)**. In addition to this isoform transition, there was also a decrease in total cTnI phosphorylation throughout postnatal development **(Figure 1H),** likely localized to residues Ser 22 and 23 **(Figure S1H)**. Overall, these data indicate that the sarcomere, specifically TnI, is dynamically regulated during the metabolic transition of glycolysis to fatty acid oxidation and during the loss of the regenerative capacity of neonatal mouse hearts.

### ssTnI overexpression does not strongly alter the global cardiac proteome in juvenile or young adult mouse hearts

Since the ssTnI to cTnI switch was one of the largest changes in protein expression detected during postnatal mouse heart development, we wanted to determine how ssTnI overexpression affects baseline cardiac protein expression in mice. To do so, we utilized the Tnni1 transgenic (Tg) mouse line driven by the alpha-myosin heavy chain (α-MHC) promoter ^13,21^ **(Figure 2A)**. Using our highly quantitative top-down proteomics platform, we were able to robustly measure TnI expression in WT and Tnni1 Tg mice **(Figure S2A-C)**. We found that Tnni1 Tg mice maintain higher levels of ssTnI at P8 when compared to WT mice, whereas cTnI expression was undetectable **(Figure 2B)**. Furthermore, P28 Tnni1 Tg mice express no detectable levels of cTnI, indicating that overexpression of ssTnI completely prevents cTnI expression in the postnatal heart **(Figure 2B)**. Interestingly, while the Tnni1 Tg mouse model has been extensively characterized before, using top-down proteomics, we were also able to identify a +190 Da mass shift in the Tg ssTnI compared to WT ssTnI **(Figure S2D)**. We were able to localize this mass shift to amino acid residues 23-82 but were not able to confirm the source of the modification. Importantly, we were not able to identify any unmodified proteoforms; thus, it is highly likely this mass shift is due to a mutation along the Tg ssTnI amino acid backbone. As this mouse model has been well characterized, we do not believe this mutation affects proteoform or heart function.

**Figure 2.**
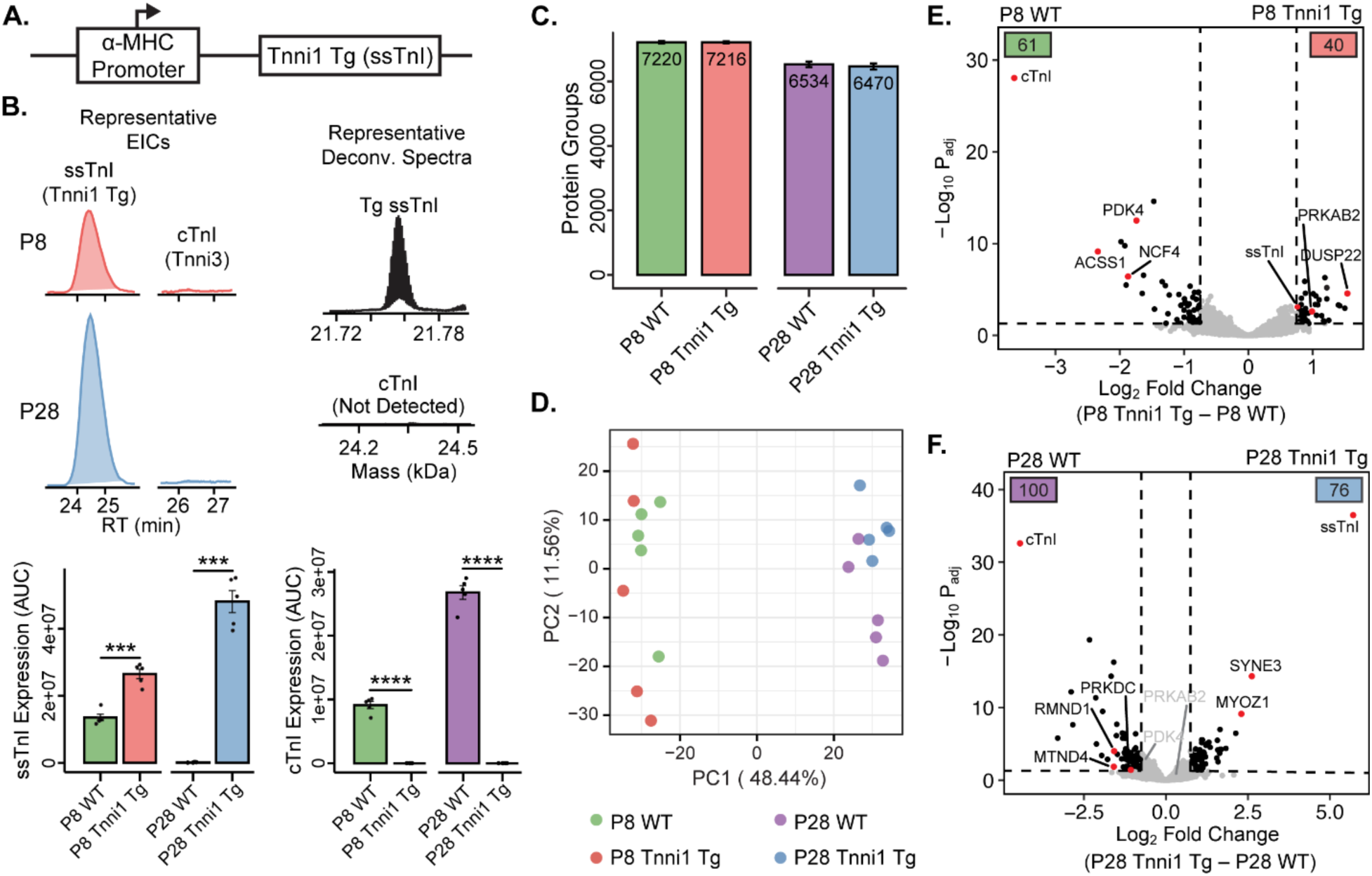
ssTnI overexpression does not strongly alter the global cardiac proteome in juvenile or adult mouse hearts. **A.** Schematic of the Tnni1 transgene. Tnni1 (ssTnI) expression is driven by the alpha-myosin heavy chain (α-MHC) promoter, extending expression of ssTnI into adulthood. **B.** Top-down proteomic analysis of ssTnI (Tnni1) and cTnI (Tnni3) expression throughout postnatal development in WT and Tnni1 Tg mice. Top: Representative EICs and deonconvoluted spectra of ssTnI (Tnni1) and cTnI (Tnni3) in the Tnni1 Tg mice. Bottom: Expression level analysis of ssTnI and cTnI in P8 and P28 WT and Tnni1 Tg mouse hearts (n = 5 per group, bars represent average, error bars represent S.E.M., dots represent individual data points. *** denotes *p* ≤ 0.001, and **** denotes *p* ≤ 0.0001 as determined by one-way ANOVA followed by Tukey’s HSD). **C.** Bar plot representing the average number of protein groups identified by global bottom-up proteomics of P8 and P28 WT and Tnni1 Tg mouse hearts demonstrating similar number of protein group identifications between the groups (n = 5 per group, error bars represent S.E.M.) **D.** Principal component analysis (PCA) of per-sample Log_2_ protein abundances displays high similarity between WT and Tnni1 Tg mice baseline proteomes at P8 and P28. **E and F.** Volcano plot demonstrating fold-change in protein expression between WT and Tnni1 Tg hearts at P8 **(E)** and P28 **(F)**. The number of significantly upregulated proteins per group is shown in the bottom corners of each comparison (n = 5 per group, adjusted p-value ≤ 0.05 and |Log2 Fold Change| ≥ 0.75 significance thresholds).

Next, using global bottom-up proteomics, we were able to quantify over 7,000 protein groups among P8 and P28 WT and Tnni1 Tg mouse hearts **(Figure 2C, Table S3)**. Quantified proteins demonstrated low CVs, providing higher confidence in our downstream analyses **(Figure S2E).** Interestingly, after performing dimensional reduction using PCA, both WT and Tnni1 Tg mice demonstrate highly similar baseline proteomes, as both WT and Tnni1 Tg groups clustered closely together at P8 and at P28 **(Figure 2D)**. To investigate the differences in proteome composition more closely, we generated volcano plots comparing WT and Tnni1 Tg mice at P8 and P28 **(Figure 2E-F)**. At P8, we detected very few DEPs, with 61 proteins upregulated in WT hearts and 40 proteins upregulated in Tnni1 Tg hearts. Unsurprisingly, the most differentially expressed protein was cTnI, which showed markedly higher expression in WT hearts **(Figure 2E and S2F)**. In WT and Tnni1 Tg mice at P8, there were changes in the expression of key metabolic proteins. Specifically, WT mice showed an increased expression of PDK4, which can reduce glycolysis utilization and inhibit cardiac regeneration^22^. On the other hand, Tnni1 Tg mice exhibited an increased expression of AMPK (PRKAB2 subunit), which is a central metabolic regulator that can modulate both glycolysis and fatty acid oxidation^23^. These changes suggest a potential shift in the baseline metabolic activities between WT and Tnni1 Tg mice. The differences in metabolic protein levels become even more pronounced when comparing the hearts of P28 WT and Tnni1 Tg mice **(Figure 2F)**. Moreover, GO analysis of the proteins upregulated in P28 WT hearts indicates the enrichment of many different biological processes related to metabolism, including oxidation-reduction processes (GO:0055114) **(Figure S2G, Table S3).** Together, these data highlight an impact of TnI isoform expression on cardiac metabolic protein expression, where ssTnI overexpression promotes a more immature cardiac metabolic state compared to WT mice.

### ssTnI overexpression promotes cardiomyocyte proliferation and cardiac regeneration in juvenile mice after myocardial infarction

As the cardiac metabolic state directly influences the regenerative potential of the heart, we next sought to determine if the altered metabolic state of the Tnni1 Tg mice affects the response to cardiac injury. To do so, we performed MI at P7, a timepoint where wild-type hearts naturally scar rather than regenerate, in both WT and Tnni1 Tg mice. Hearts were then harvested at 7 days post-surgery (DPS) to assess CM proliferation and 21 DPS to assess scar formation and CM size **(Figure 3A)**. Interestingly, there was a significant increase in the number of CMs undergoing mitosis as demonstrated by phospho-histone H3 (pH3) staining in Tnni1 Tg mice compared to WT mice at 7 days post-MI at P7 **(Figure 3B)**. Additionally, there was a significant increase in the number of CMs undergoing cytokinesis by Aurora B staining in the Tnni1 Tg mice compared to WT mice **(Figure 3C)**. These results indicate that Tnni1 Tg CMs display increased mitosis and cytokinesis following injury, suggesting that more CMs undergo complete cell division following ssTnI overexpression. At 21 DPS, there was no difference in CM cross-sectional area in sham-operated animals as quantified by wheat germ agglutinin staining (WGA) **(Figure S3A)**; however, WT CMs were significantly larger than Tnni1 Tg CMs post-MI **(Figure 3D)**, which is consistent with reduced hypertrophy and a higher number of newly formed smaller CMs in Tnni Tg mice. Lastly, to determine whether the increased CM proliferation in enhanced the regenerative response of Tnni1 Tg mice, we performed trichrome staining to quantify scar size and myocardial regeneration. Remarkably, we quantified a significantly smaller scar size in Tnni1 Tg mice compared to the large fibrotic scars in WT mice at 21 DPS **(Figures 3F and S3D-E).** Taken together, these findings indicate that replacement of cTnI with ssTnI in the postnatal heart promotes robust cardiac regeneration after P7 MI and this regenerative process is driven by CM proliferation.

**Figure 3.**
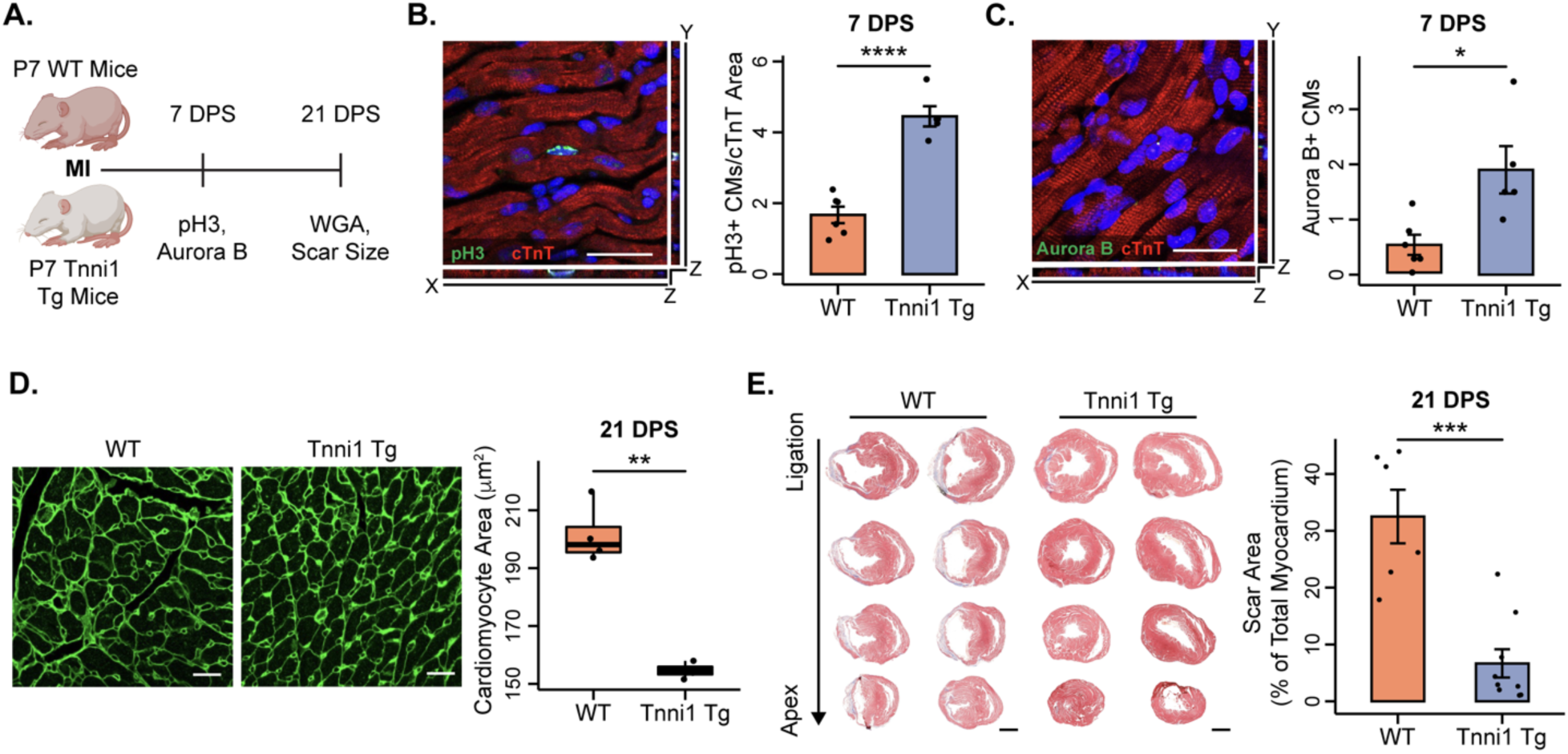
ssTnI overexpression promotes cardiomyocyte proliferation and cardiac regeneration in juvenile mice after myocardial infarction. **A.** Workflow of experimental design. Myocardial infarction surgeries were performed on P7 WT (non-regenerative) and P7 Tnni1 Tg mice. 7 days post-surgery (DPS), hearts were harvested to assess cardiomyocyte proliferation. 21 DPS, hearts were harvested to assess cardiomyocyte size and scar formation. **B.** Left: High magnification Z-stack image of a mitotic cardiomyocyte after immunostaining phosphohistone H3 staining (pH3) and cardiac troponin T (cTnT) at 7 DP P7 MI. Scale bar, 25 µm. Right: Quantification of the number of mitotic cardiomyocytes normalized by cTnT area. P7 MI Tnni1 Tg hearts display a significant increase in cardiomyocyte proliferation 7 DPS (n = 6 WT hearts, n = 5 Tnni1 Tg hearts, 4 sections quantified per heart, bars represent average, error bars represent S.E.M., dots represent average per heart). **C.** Left: High magnification Z-stack image of a cardiomyocyte going through cytokinesis after immunostaining for Aurora B kinase and cTnT 7 DP P7 MI. Scale bar, 25 µm. Right: Quantification of the number of cardiomyocytes going through cytokinesis. P7 MI Tnni1 Tg hearts display a significant increase in cardiomyocyte cytokinesis 7 DPS (n = 6 WT hearts, n = 5 Tnni1 Tg hearts, 4 sections quantified per heart, bars represent average, error bars represent S.E.M., dots represent average per heart). **D.** Left: Wheat germ agglutinin (WGA) staining at 21 DP P7 MI to assess cardiomyocyte size. Scale bar, 20 µm. Right: Quantification of cardiomyocyte area by WGA staining. P7 MI Tnni1 Tg cardiomyocytes are significantly smaller than WT cardiomyocytes 21 DPS (n = 4 hearts per group, n = 3 sections per heart, n = 150 cardiomyocytes per section. Boxplots display median, upper, and lower quartile; dots represent average cardiomyocyte area per heart). **E.** Left: Masson’s trichrome staining of WT and Tnni1 Tg mice hearts 21 DP P7 MI. Right: Quantification of scar area as a percent of total viable myocardium. P7 MI Tnni1 Tg hearts display a significant decrease in scar formation compared to WT hearts 21 DPS (n = 6 WT hearts, n = 9 Tnni1 Tg hearts, 3 sections quantified per heart, bars represent average, error bars represent S.E.M., dots represent average per heart). * denotes *p* ≤ 0.05, ** denotes *p* ≤ 0.01, *** denotes *p* ≤ 0.001, and **** denotes *p* ≤ 0.0001 as determined by a 2-tailed unpaired Student’s *t*-test.

### ssTnI overexpression promotes elevated expression of AMPK, glycolytic enzymes, and proteasomal subunits during regeneration

To understand the molecular basis of the prolonged regenerative capacity of Tnni1 Tg mice, we performed MI on P1 WT (regenerative), P7 WT (non-regenerative) and P7 Tnni1 Tg (regenerative) mice, collected the hearts at 7 DPS, and analyzed the ventricular tissue using global bottom-up and targeted sarcomeric top-down proteomics **(Figure 4A)**. In our global analysis, we quantified over 7,000 protein groups with low CVs **(Figure S4A-B, Table S5)**. Additionally, when we applied PCA to reduce the dimensionality of the data, it revealed three distinct clusters, each corresponding to a specific injury response **(Figure 4B)**. Interestingly, when condensed along the y-axis (PC2), the two regenerative responses (P1 WT and P7 Tnni1 Tg) clustered closely, indicating a shared proteomic response to injury. Interestingly, a hierarchical heatmap of the DEPs between the three conditions demonstrates three distinct patterns of protein expression in response to injury **(Figure 4C)**.

**Figure 4.**
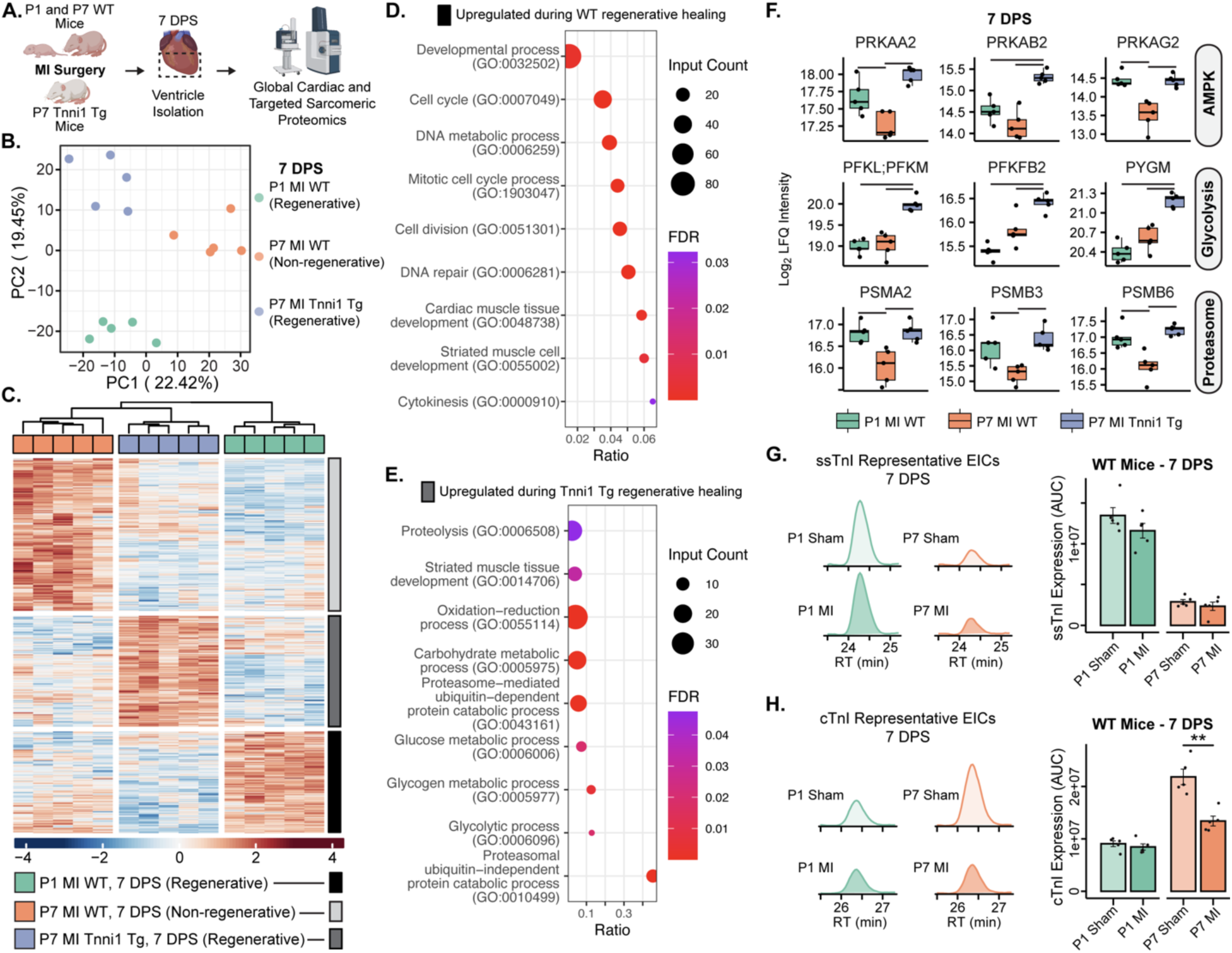
ssTnI overexpression induces levels of AMPK, glycolytic enzymes, and proteasomal subunits after myocardial infarction compared to non-regenerative WT mice. **A.** Workflow of experimental design. Myocardial infarction surgeries were performed on P1 WT (regenerative), P7 WT (non-regenerative) and P7 Tnni1 Tg mice (regenerative) (n = 5 per group). 7 days post-surgery (DPS), hearts were harvested for global bottom-up cardiac proteomics and targeted top-down sarcomeric proteomics. **B.** Principal component analysis (PCA) of per-sample Log_2_ protein abundances displays three distinct responses to injury. Regenerative responses to injury cluster closely along PC1, while P7 injuries cluster closely along PC2. **C.** Hierarchal heatmap (k-means columns set to 3, k-means rows set to 3) displaying z-score normalized protein intensities of all significantly differentially expressed proteins throughout postnatal development (adjusted p-value ≤ 0.05 and |Log2 Fold Change| ≥ 0.75 significance thresholds). The rows are separated into three clusters: proteins with elevated expression in during a non-regenerative response to injury (light grey), proteins with elevated expression during a WT regenerative response to injury (black), and proteins elevated expression during a Tnni1 Tg regenerative response to injury (dark grey). **D and E.** Selected STRING biological process gene ontology (GO) plots of the proteins that are elevated during WT regenerative healing **(D)** and Tnni1 Tg regenerative healing **(E)**. GO analysis for WT non-regenerative healing is included in Figure S8. Ratio represents the fractions of all proteins in the GO category that were identified. Dot size corresponds to the number of identified proteins within a GO category. Color represents FDR-adjusted p-value of the overrepresentation test. **F.** Boxplots of selected proteins related to AMPK, glycolysis, and the proteasome upregulated during Tnni1 Tg regenerative response to injury. (n = 5 per group, boxplots display median, upper, and lower quartile; dots represent individual hearts; bars between groups represent an adjusted p-value ≤ 0.05). **G and H.** Top-down proteomic analysis of ssTnI expression **(G)** and cTnI expression **(H)** 7 DP P1 or P7 sham or MI surgeries. Left: Representative EICs of ssTnI and cTnI 7 DP P1 or P7 sham or MI surgeries. Right: Expression level analysis of ssTnI and cTnI 7 DP P1 or P7 MI. No change detected in ssTnI expression during regenerative (P1) or non-regenerative response (P7) to MI. Decreased expression of cTnI detected during non-regenerative (P7) response to MI, indicating cTnI expression may be regulated post-injury (n = 5 per group, bars represent average, error bars represent S.E.M., dots represent individual data points. ** denotes *p* ≤ 0.01 as determined by a 2-tailed unpaired Student’s *t*-test).

When using STRING to conduct a GO analysis, the P7 MI WT samples displayed a typical non-regenerative injury response, which included apoptotic cell clearance (GO:0043277), activation of immune response (GO:0002253), and fibrinolysis (GO:0042730) (**Figure S4C, Table S5)**. Conversely, the GO terms enriched for the P1 MI WT samples were more in line with regenerative injury response, such as mitotic cell cycle process (GO:1903047), cardiac muscle tissue development (GO:0048738), and cytokinesis (GO:0000910) **(Figure 4D, Table S5)**. On the other hand, GO analysis of the P7 MI Tnni1 Tg samples highlighted a distinct regenerative response to injury governed by increased proteolytic processes (GO:0006508, GO:0043161, and GO:0010499) and glycolytic metabolic processes (GO:0005975, GO:0006006, GO:0005977) **(Figure 4E, Table S5)**. These results suggest that the increased CM proliferation of the Tnni1 Tg mice may be attributed to increased glycolytic metabolism, a known regulator of CM proliferation.

Of note, previous studies demonstrated that ssTnI overexpression provided protection against ischemia-reperfusion^14^ and chronic pressure-overload^15^. This protection was attributed to increased AMPK activity and glycolytic metabolism. Congruently, our results reveal a pronounced increase in all subunits of both the cardiac (PRKAA2, PRKAB2, and PRKAG2) and general (PRKAA1, PRKAB1, PRKAG1) AMPK isoforms (Figures 4F, S4D). Additionally, there was an elevated expression of specific downstream glycolytic enzymes, notably PFKL, PFKM, and PGKFB2, which are under direct regulation by AMPK **(Figures 4F, S4E)**. Together, our results emphasize the intricate relationship between the sarcomere, metabolic regulation, and cellular proliferation. In addition to upregulation of AMPK and glycolytic proteins, we also observed a significant increase in levels of many subunits of the 20S proteasome (PSMA2, PSMA3, PSMA7, PSMB1, PSMB2, PSMB3, PSMB4, PSMB5, and PSMB6) **(Figures 4F, S4F)**. Recent studies indicate the proteasome’s important role in promoting CM proliferation during natural cardiac regeneration^24^. Moreover, higher levels of the 20S proteasome have been linked to enhanced cell survival during conditions of hypoxia and cellular stress^25^.

Next, we compared P1 MI and P7 MI WT mice at 7 DPS to sham animals to determine whether ssTnI or cTnI expression is differentially regulated during regeneration or pathological remodeling. From our comprehensive proteomic analysis of the cardiac sarcomere, we found that ssTnI expression remained unchanged after both P1 and P7 MI (**Figure 4G**). Interestingly, while the phosphorylation levels of cTnI were consistent (**Figure S4G)**, there was a marked reduction in cTnI abundance in WT mice at 7 DPS following a P7 MI compared to sham controls (**Figure 4H**). This decrease in cTnI expression may be a protective mechanism following injury as the hearts are attempting to promote CM dedifferentiation and subsequent CM proliferation ^26^.

Lastly, we performed a global proteomic analysis of ventricular tissue from P7 Tnni1 Tg mice 7 days-post MI or sham surgery to understand how injury alters proteome composition **(Figure S5A)**. When examining the DEPs after sham or MI surgery in the Tnni1 Tg mice **(Figure S5B)**, there was a significant downregulation of proteins related to oxidative phosphorylation during regeneration in Tnni1 Tg mice **(Figure S5C),** while there was an upregulation of immune system processes (GO:0002376) and regeneration processes (GO:0031099) during regeneration in Tnni1 Tg mice **(Figure S5D)**. Interestingly, many subunits of both the general and cardiac specific isoforms of AMPK were not differentially expressed between sham and MI Tnni1 Tg mice **(Figure S5E)**, indicating that the elevated AMPK expression is a baseline phenotype of the Tnni1 Tg mice that may provide the metabolic flexibility necessary for CM proliferation after injury. Unlike AMPK, there was significant downregulation of proteins related to oxidative-phosphorylation (PDK4, NDUFS3, and ATP5PO) and significant upregulation of proteins involved in the proteasome and cell proliferation following injury **(Figure S5F).** These data indicate that the regenerative injury response of Tnni1 Tg mice is likely due to an increased metabolic flexibility augmented by a baseline increase in AMPK expression.

Collectively, our results reveal a dynamic interplay between sarcomere isoforms and cardiac metabolism, which uncovers a molecular mechanism that can be targeted to control cardiac metabolism, sarcomere disassembly, CM proliferation, and heart regeneration.

## DISCUSSION

One major barrier to CM proliferation is the large size of the sarcomeres, the contractile proteins of CMs, which are not conducive to cell division^27,28^. Thus, CMs need to disassemble their sarcomeres for proliferation to proceed during heart regeneration^8,29^. The mechanisms that control sarcomere disassembly and reassembly are not well understood. Importantly, rewiring adult cardiac metabolism can induce sarcomere disassembly, CM proliferation, and heart regeneration^16,18^. However, the interplay between sarcomeres and cardiac metabolism remains undefined.

Interestingly, a sarcomere isoform switch from ssTnI to cTnI takes place postnatally, which is concomitant with the metabolic switch that contributes to loss of the cardiac regenerative potential. We wanted to determine whether this sarcomere switch impacts cardiac metabolism. In this study, we dissect the proteomic and sarcomere transitions during postnatal maturation by quantitative bottom-up and top-down proteomics. Importantly, we demonstrate that ssTnI overexpression in the postnatal heart increases the cardiac regenerative capacity following injury, displaying a significant increase in CM proliferation and reduction in scar size. We conducted a global quantitative proteomic analysis of ssTnI transgenic mice after injury, which demonstrates that the overexpression of ssTnI shifts the cardiac metabolic landscape towards glycolysis following injury. Our results reveal that ssTnI regulates a sarcomere-metabolism axis that is required for CM proliferation and heart regeneration.

Elucidating the molecular mechanisms by which sarcomere isoforms control metabolic processes could offer key insights into regulating both cardiac metabolism and regeneration. Interestingly, naked mole-rats, known for their remarkable tolerance to extreme hypoxia and anoxia due to metabolic adaptability, display a distinctive myofilament protein composition in their cardiac ventricles that includes the co-expression of cTnI and ssTnI^30,31^. Yet, it remains undetermined whether adult naked mole-rats possess the ability to regenerate their hearts after injury. Furthermore, recent studies suggest a potential role for Ca^2+^ signaling in regulating CM proliferation and cardiac regeneration^32,33^. It is unclear whether the modulation of sarcomere sensitivity to Ca^2+^ plays a pivotal role during heart regeneration. Future studies are required to define this cross talk between sarcomeres and metabolism by biochemical, and metabolomic approaches to provide a framework for metabolic control of sarcomere disassembly and CM proliferation. Our study uncovers a novel role in the interplay between sarcomeres and cardiac metabolism, representing a potential therapeutic target to overcome a significant obstacle in CM proliferation and regeneration.

## MATERIALS AND METHODS

### Reagents and Chemicals

All reagents were purchased from Sigma-Aldrich, Inc. (St. Louis, MO, USA) unless otherwise noted. High performance mass spectrometry (MS)-grade water, acetonitrile, and ethanol were purchased from Fischer Scientific (Fair Lawn, NJ, USA).

### Animals

CD-1 mice were procured from Charles River Laboratories. Tnni1 Tg mice were obtained from R. John Solaro and Beata M. Wolska and were maintained at the University of Wisconsin-Madison. All experimental procedures performed on animals were approved by the Institutional Animal Care and Use Committee of the University of Wisconsin-Madison. All experiments were performed on age-matched mice, with a roughly equal ratio of male to female mice for neonatal experiments.

### Neonatal Myocardial Infarction (MI)

Surgeries were performed on neonatal mice at postnatal day (P) 1 or P7. Sham and MI surgeries were performed as previously described ^34^. In brief, neonatal mice were anesthetized by hypothermia on ice. After skin incision, a lateral thoracotomy at the fourth intercostal space was performed by dissection of the intercostal muscles. The heart was exposed by applying pressure to the chest, and a C-1 tapered needle attached to a 6-0 Prolene suture (Ethicon Inc, Bridgewater, NJ, USA Cat# EPM8706) was passed through the ventricle below the left anterior descending coronary artery and tied off to induce MI. The ribs and chest wall were closed using the same suture, and the skin was closed with adhesive glue (3M, St. Paul, MN, USA Cat# 1469SB). The neonates were warmed on a heating pad until recovered. Sham-operated mice underwent the same procedure, but the left anterior descending coronary artery was not ligated.

### Immunofluorescent Staining

Following dissection, the atria were removed from the hearts and briefly washed in PBS to remove blood. The hearts were then fixed using 4% paraformaldehyde (PFA) in PBS overnight at 4 ℃, washed with PBS, and then dehydrated in 70% ethanol overnight before paraffin embedding. 5 µM serial sections below the site of ligation were placed onto glass slides, and the tissue was then deparaffinized, rehydrated, and boiled in IHC antigen retrieval solution (Invitrogen, Waltham, MA, USA Cat# 00-495558) for 10 min in a microwave. Sections were blocked with 5% goat serum (Vector Laboratories, Newark, CA, USA, Cat# S-1000) in PBS, and then incubated overnight at 4 ℃ with primary antibodies diluted in 5% goat serum in PBS: phospho-histone H3 Ser10 (Millipore, Burlington, MA, USA, Cat# 06-570, 1:100 dilution), aurora B Kinase (Sigma, St. Louis, MO, USA Cat# A5102, 1:100 dilution), and cardiac troponin T (Abcam, Cambridge, UK, Cat# ab829, 1:100 dilution). Sections were then washed PBS and then incubated for 1 h at room temperature with secondary antibody solution containing 5% goat serum in PBS and the appropriate secondary antibodies conjugated to Mouse-555 (Invitrogen, Waltham, MA, USA, Cat# A28180, 1:400 dilution) or Rabbit-488 (Invitrogen, Waltham, MA, USA, Cat# A-11008, 1:400 dilution). Nuclei were than stained using DAPI, and the slides were mounted using Fluoromount-G (ThermoFisher, Waltham, MA, USA, Cat# 00-4958-02). Slides were imaged on a Keyence BZ-X800 microscope, and high magnification images were captured on Nikon A1RS HD confocal microscope. Raw values and statistical analysis are available in **Table S4**.

### Cardiomyocyte Cross-Sectional Area

Paraffin-embedded hearts were serially sectioned in 5 µM slices below the site of ligation onto glass slides. The sections were deparaffinized, rehydrated, and boiled in IHC antigen retrieval solution (Invitrogen, Waltham, MA, USA, Cat# 00-495558) for 10 min in a microwave. Sections were stained with a wheat germ agglutinin (WGA) antibody that was pre-conjugated with Alexa Fluor 488 (ThermoFisher Scientific, Waltham, MA, USA, Cat# W11261) for 1 h at room temperature, washed with PBS-Tween (0.5%), and then mounted using Fluoromount-G (ThermoFisher Scientific, Waltham, MA, USA, Cat# 00-4958-02). Slides were imaged on a Keyence BZ-X800 microscope, and high magnification images were captured on Nikon A1RS HD confocal microscope. ImageJ was used to quantify cardiomyocytes by measuring the cross-sectional area of 150 cardiomyocytes per section, with at least 3 sections measured per heart. Average cardiomyocyte area for each heart was plotted. Raw values and statistical analysis are available in **Table S4**.

### Histology

Paraffin-embedded hearts were serially sectioned in 5 µM slices below the site of ligation onto glass slides. Masson trichrome staining was performed following the manufacturer’s protocol (Newcomer Supply, Middleton, WI, USA, Cat# 9179). Scar size measurements were quantified from at least 3 sections from each heart. Fibrotic scar was quantified using ImageJ, and the average scar area for each heart was plotted. Raw values and statistical analysis are available in **Table S4**.

### Global Quantitative Proteomic Analysis

To create an atlas of postnatal cardiac development, Tnni1 Tg mice, and myocardial infarction injury response, we collected hearts (n = 5 per group) from mice as outlined in **Table S1**. Hearts were removed from the mice, the atria were dissected away from the heart, and then the tissues were quickly washed in PBS to remove blood. Ventricular tissue was snap-frozen in liquid nitrogen and then cryopulverized.

With minor modifications, the global ventricular proteome was extracted as previously described ^19,35^. In a cold room (4 ℃), samples were homogenized using a Teflon pestle in 0.2% Azo lysis buffer (0.2% w/v Azo, 25 mM ammonium bicarbonate (ABC), 10 mM L-methionine, 1 mM DTT, and 1X HALT protease and phosphatase inhibitor (ThermoFisher Scientific, Waltham, MA, USA Cat# 78440)). Samples were further homogenized in a sonicating water bath for 10 min at 4 ℃ prior to centrifugation at 21,100 *g* for 30 min at 4 ℃. The cleared supernatant was transferred to a new tube, and aliquots were 1:50 in water prior to Bradford protein assay (Bio-Rad, Hercules, CA, USA, Cat# 5000006). Samples were then normalized to 1 mg/mL protein in 0.1% Azo prior to reduction with 30 mM DTT at 37C for 1 h and alkylation with 30 mM chloroacetamide for 45 min. Following reduction and alkylation, samples were digested overnight on a 1000 rpm shaker at 37 ℃ with Trypsin Gold (Promega, Madison, WI, USA, Cat# V5820) in a 50:1 ratio (wt/wt) of protein:protease. The reaction was halted with the addition of 1% FA, Azo was degraded by applying a 305 nm UV lamp (UVN-57 Handheld UV Lamp; Analytik Jena, Jena, TH, DEU) for 5 min, and then samples were cleared at 21,100 g at 4 °C for 30 min. Peptides were desalted using 100 µL Pierce C18 tips (ThermoFisher Scientific, Waltham, MA, USA, Cat#87784) using the manufacturer’s protocol and then dried in a vacuum centrifuge prior to resuspension in 0.1% FA. Peptide concentrations were measured using A205 readings on a NanoDrop.

200 ng protein digest was separated by reverse-phase liquid chromatography using a nanoElute nano-flow ultra-high pressure LC system (Bruker Daltonics) and analyzed on a trapped ion mobility quadrupole-time-of-flight mass spectrometer (timsTOF Pro, Bruker Daltonics) following CaptiveSpray nano-electrospray ionization. Briefly, peptides were washed on a PepMap Neo C18 trap column (ThermoFisher Scientific, Waltham, MA, USA, Cat#174500) for 10-min at 2% mobile phase B (mobile phase A (MPA): 0.1% FA; mobile phase B (MPB): 0.1% FA in acetonitrile), and then separated on a C18 analytical column (25 cm length, 75 µm inner diameter, 1.6 µm particle size, 120 Å pore size; IonOpticks, Fitzroy, VIC, AUS, Cat# 1801220) heated to 55 ℃ at a flow rate of 400 nL/min using a gradient of 2-17% MPB from 10-70 min, 17-25% MPB from 70-100 min, 25-37% MPC from 100-110 min, and 37-85% MPB from 110-120 min, followed by a 10 min wash at 85% MPB. Samples were collected in data-indepdent analysis-parallel accumulation-serial fragmentation (diaPASEF) mode mode using 32 windows ranging from 0.6 to 1.421/K 0 and 400 to 1200 m/z^36^, as previously described.

All raw mass spectrometry (MS) files were processed together with DIA-NN in double-pass mode using an *in silico* generated spectral library from the mouse Uniprot proteome (UP000000589, reviewed, no isoforms, accessed 20 April 2022) ^37^. The following parameters were employed: peptide length required to be between 200 and 1700 *m/z*, no more than 2 missed cleavages allowed, cysteine carbamidomethylation was set as a fixed modification, methionine oxidation and N-terminal acetylation were set as variable modifications, mass accuracy was set to 15 ppm, heuristic protein grouping was turned off, match-between-runs was enabled at 1% false-discovery rate (FDR), and heuristic protein grouping was turned off. Protein and peptide output is included in **Table S1**.

Comparative analyses were performed in R (version 4.1.0.) using DAPAR ^38^, DEP ^39^ and IHW ^40^ as previously described. First, samples were filtered to remove proteins that were not quantified in at least three of five biological replicates within one sample group. Then, quantification values were Log_2_-transformed and normalized to the median of the total data set. Partially observed values within a sample group were imputed via ssla, and values missing across an entire sample group were imputed at the 2.5% quantile. Limma tests were utilized to evaluate statistical significance, and *p-values* were adjusted via independent hypothesis weighting based on the number of peptides observed per protein group. For proteome wide comparisons (volcano plots, heatmaps, and gene ontology analyses), statistical significance required a Log2-fold change of 0.75 or greater in either direction and an FDR-adjusted p-value of 0.05 or less. For individual comparisons (boxplots), statistical significance required an FDR-adjusted p-value of 0.05 or less. Gene ontology (GO) analyses were conducted using STRING, requiring an FDR-adjusted *p-value* of 0.05 for a GO term to be considered statistical enriched. A list of differentially expressed proteins and GO analysis results are included in **Tables S2, S3, and S5**.

### Top-down Sarcomere Proteomic Analysis

Using the same samples (n = 5 per group) detailed in **Table S1**, sarcomeric proteins were extracted as previously described with minor modification ^41^. Briefly, in a cold room (4 ℃) ventricular tissue was homogenized with a Teflon pestle (1.5 mL micro-centrifuge tube flat-tip; Thomas Scientific, Swedesboro, NJ, USA) in 100 µL HEPES buffer (25 mM HEPES, pH 7.4, 60 mM NaF, 10 mM L-Methionine, 1 mM DTT, 1 mM PMSF, 1 mM Na_3_VO_4_, and 1X HALT protease and phosphate inhibitor cocktail (ThermoFisher Scientific, Waltham, MA, USA, Cat# 78440)). The samples were centrifuged, and the pellet was homogenized in another 100 µL HEPES buffer to fully deplete cytosolic proteins. The homogenate was centrifuged at 21,100 *g* for 30 min at 4 ℃, and the remaining pellet was transferred to a new tube and homogenized in 50 µL trifluoroacetic acid (TFA) solution (1% TFA, 1 mM TCEP, 10 mM L-Methionine). The sarcomere-enriched extracts were cleared by centrifugation, and the resulting supernatant was diluted 10-fold prior to Pierce™ 660 nm Protein Assay (ThermoFisher Scientific, Waltham, MA, USA, Cat# 22660). The samples were normalized to 100 ng/µL in 0.1% FA with 2 mM TCEP for LC-MS analysis. Additional samples were prepared at 25, 50, 150, and 200 ng/µL to generate linear curves.

Mass spectra were acquired on a high-resolution Impact II quadrupole time-of-flight (Q-TOF) mass spectrometer (Bruker Daltonics, Bremen, Germany) coupled to a NanoAcquity ultra-high pressure LC system (Waters, Milford, MA, USA). After 1 min of washing at 25 µL/min and 95% MPA (MPA: 0.1% formic acid (FA) in water; MPB: 0.1% FA in 50:50 acetonitrile:ethanol), 500 ng sarcomeric protein was separated on a home-packed PLRP column (PLRP-S, 100 Å pore size, 10 µm particle size, 500 µm inner diameter) by reverse-phase liquid chromatography (RPLC) using a stepwise gradient of 5-20% MPB from 10-15 min, 20-40% MPB from 15-27 min, 40-50% MPB from 27-35 min, 50-65% MPB from 35-42 min, 65-95% MPB from 42-53 min, 95% MPB from 53-58 min, then 5% MPB from 58-70 min at a constant flow rate of 12 µL/min to acquire intact protein spectra. Proteins were ionized using electrospray ionization, with the following parameters: endplate offset, 500 V; capillary voltage, 4500 V; nebulizer pressure 0.5 Bar; dry gas flow rate 4.0 L/min, and in-source collisional energy, 20 V. Mass spectra were collected from 200-3000 *m/z* at 0.5 Hz. MS/MS was performed by collision-activated dissociation (CAD) and collected at a scan rate of 1 Hz from 200-3000 *m/z*. The isolation window for online AutoMS/MS CAD ranged from 4 to 8 *m/z*. The collision DC bias was set from 18 to 45 eV for CAD with nitrogen as the collision gas.

Intact and top-down mass spectrometry data were analyzed Bruker DataAnalysis (v4.3) and Mash Native ^42^. First, chromatograms were smoothed using the Gauss algorithm (smoothing width 2.024). Extracted ion chromatograms (EICs) were created from the 3-5 most abundant charge state ions for all proteoforms in each protein family of interest, smoothed using the Gauss algorithm, and then integrated to calculate abundance. To generate deconvoluted spectra, mass spectra were averaged over the retention time (RT) window where the proteoform of interest eluted and deconvoluted using the Maximum Entropy Deconvolution algorithm (50,000 Da resolving power). As separations were consistent, the same RT window as used for all samples. MS/MS data were output from the DataAnalysis software and analyzed using MASH Native ^42^ for proteoform identification and sequence mapping. All fragment ions were manually validated with a mass tolerance of 20 ppm. Fragment ions, ions for quantification, and theoretical and experimental masses are reported in **Tables S2, S3, and S5**. Relative phosphorylation of a protein was determined using **Equation 1** as previously described ^19^, where the SNAP algorithm was used to determine the relative intensity of all proteoforms in a protein family.

**Equation 1**: Rel. Phosphorylation = (P_mono-p_ + 2 x P_bis-p_)/Total Proteoform Intensity Statistical analysis was performed using “rstatix” ^43^ in R (v4.1.0) using Student’s T-tests or one-way ANOVA followed by Tukey’s HSD post-hoc test. Data were visualized using R (version 4.1.0) with “ggplot2” ^44^ and “ggpubr” ^45^. Statistical analyses are available in **Tables S2, S3, and S5**.

## ACKNOWLEDGEMENTS

A.I.M would like to acknowledge the NIH/NHLBI HL155617, HL166256, and the DOD W81XWH2210094. Y.G. would like to acknowledge R01 HL096971, R01 GM117058, GM125085, HL109810 and S10 OD018475. T.JA. would like to acknowledge support from the Training Program in Molecular and Cellular Pharmacology, T32 GM008688 and the NIH/NHLBI under Ruth L. Kirschstein NRSA F31HL167328. W.G.P. would like to acknowledge the NIH/NHLBI under Ruth L. Kirschstein NRSA T32 HL007936 to the UW Cardiovascular Research Center.

## AUTHOR CONTRIBUTIONS

T.J.A., J.B., and W.G.P. performed the experiments and prepared the manuscript. T.J.A, J.B., Y.G., and A.I.M. designed the experiments, performed the analyses, and wrote the manuscript. E.A.C., R.J.S., and M.W.M contributed to sample processing and analysis. Y.G. oversaw proteomic experiments and wrote the manuscript. A.I.M conceived the project and wrote the manuscript.

## References

1 Tsao, C. W. et al. Heart Disease and Stroke Statistics—2023 Update: A Report From the American Heart Association. Circulation 147, e93–e621, doi:doi:10.1161/CIR.0000000000001123 (2023).

2 Murphy, S. P., Ibrahim, N. E. & Januzzi, J. L., Jr. Heart Failure With Reduced Ejection Fraction: A Review. JAMA 324, 488–504, doi:10.1001/jama.2020.10262 (2020).

3 Porrello, E. R. et al. Transient Regenerative Potential of the Neonatal Mouse Heart. Science 331, 1078–1080, doi:10.1126/science.1200708 (2011).

4 Mahmoud, A. I., Porrello, E. R., Kimura, W., Olson, E. N. & Sadek, H. A. Surgical models for cardiac regeneration in neonatal mice. Nature Protocols 9, 305–311, doi:10.1038/nprot.2014.021 (2014).

5 Siedner, S. et al. Developmental changes in contractility and sarcomeric proteins from the early embryonic to the adult stage in the mouse heart. The Journal of Physiology 548, 493–505, 10.1111/j.1469-7793.2003.00493.x (2003).

6 Guo, Y. & Pu, W. T. Cardiomyocyte Maturation. Circulation Research 126, 1086–1106, doi:10.1161/CIRCRESAHA.119.315862 (2020).

7 Mahmoud, A. I. et al. Meis1 regulates postnatal cardiomyocyte cell cycle arrest. Nature 497, 249–253, doi:10.1038/nature12054 (2013).

8 Porrello, E. R. et al. Transient regenerative potential of the neonatal mouse heart. Science 331, 1078–1080, doi:10.1126/science.1200708 (2011).

9 Bae, J., Paltzer, W. G. & Mahmoud, A. I. The Role of Metabolism in Heart Failure and Regeneration. Frontiers in Cardiovascular Medicine 8, doi:10.3389/fcvm.2021.702920 (2021).

10 Puente, B. N. et al. The oxygen-rich postnatal environment induces cardiomyocyte cell-cycle arrest through DNA damage response. Cell 157, 565–579, doi:10.1016/j.cell.2014.03.032 (2014).

11 Yin, Z., Ren, J. & Guo, W. Sarcomeric protein isoform transitions in cardiac muscle: A journey to heart failure. Biochimica et Biophysica Acta (BBA) - Molecular Basis of Disease 1852, 47–52, 10.1016/j.bbadis.2014.11.003 (2015).

12 Cheng, Y. & Regnier, M. Cardiac troponin structure-function and the influence of hypertrophic cardiomyopathy associated mutations on modulation of contractility. Arch Biochem Biophys 601, 11–21, doi:10.1016/j.abb.2016.02.004 (2016).

13 Fentzke, R. C. et al. Impaired cardiomyocyte relaxation and diastolic function in transgenic mice expressing slow skeletal troponin I in the heart. The Journal of Physiology 517, 143–157, 10.1111/j.1469-7793.1999.0143z.x (1999).

14 Pound, K. M. et al. Expression of slow skeletal TnI in adult mouse hearts confers metabolic protection to ischemia. Journal of Molecular and Cellular Cardiology 51, 236–243, 10.1016/j.yjmcc.2011.05.014 (2011).

15 Carley, A. N., Taglieri, D. M., Bi, J., Solaro, R. J. & Lewandowski, E. D. Metabolic Efficiency Promotes Protection From Pressure Overload in Hearts Expressing Slow Skeletal Troponin I. Circulation: Heart Failure 8, 119–127, doi:10.1161/CIRCHEARTFAILURE.114.001496 (2015).

16 Bae, J. et al. Malonate Promotes Adult Cardiomyocyte Proliferation and Heart Regeneration. Circulation, doi:10.1161/CIRCULATIONAHA.120.049952 (2021).

17 Cardoso, A. C. et al. Mitochondrial Substrate Utilization Regulates Cardiomyocyte Cell Cycle Progression. Nat Metab 2, 167–178 (2020).

18 Li, X. et al. Inhibition of fatty acid oxidation enables heart regeneration in adult mice. Nature, doi:10.1038/s41586-023-06585-5 (2023).

19 Aballo, T. J. et al. Integrated proteomics reveals alterations in sarcomere composition and developmental processes during postnatal swine heart development. Journal of Molecular and Cellular Cardiology 176, 33–40, 10.1016/j.yjmcc.2023.01.004 (2023).

20 Bae, J. et al. Malonate Promotes Adult Cardiomyocyte Proliferation and Heart Regeneration. Circulation 143, 1973–1986, doi:10.1161/CIRCULATIONAHA.120.049952 (2021).

21 Wolska, B. M. et al. Expression of slow skeletal troponin I in adult transgenic mouse heart muscle reduces the force decline observed during acidic conditions. The Journal of Physiology 536, 863–870, 10.1111/j.1469-7793.2001.00863.x (2001).

22 Cardoso, A. C. et al. Mitochondrial substrate utilization regulates cardiomyocyte cell-cycle progression. Nature Metabolism 2, 167–178, doi:10.1038/s42255-020-0169-x (2020).

23 Arad, M., Seidman, C. E. & Seidman, J. G. AMP-Activated Protein Kinase in the Heart. Circulation Research 100, 474–488, doi:10.1161/01.RES.0000258446.23525.37 (2007).

24 Cui, M. et al. Nrf1 promotes heart regeneration and repair by regulating proteostasis and redox balance. Nature Communications 12, 5270, doi:10.1038/s41467-021-25653-w (2021).

25 Sahu, I. et al. The 20S as a stand-alone proteasome in cells can degrade the ubiquitin tag. Nature Communications 12, 6173, doi:10.1038/s41467-021-26427-0 (2021).

26 Wang, W. E. et al. Dedifferentiation, Proliferation, and Redifferentiation of Adult Mammalian Cardiomyocytes After Ischemic Injury. Circulation 136, 834–848, doi:10.1161/CIRCULATIONAHA.116.024307 (2017).

27 Ahuja, P., Perriard, E., Perriard, J. C. & Ehler, E. Sequential myofibrillar breakdown accompanies mitotic division of mammalian cardiomyocytes. J Cell Sci 117, 3295–3306, doi:10.1242/jcs.01159 (2004).

28 Kubin, T. et al. Oncostatin M is a major mediator of cardiomyocyte dedifferentiation and remodeling. Cell Stem Cell 9, 420–432, doi:10.1016/j.stem.2011.08.013 (2011).

29 Jopling, C. et al. Zebrafish heart regeneration occurs by cardiomyocyte dedifferentiation and proliferation. Nature 464, 606–609, doi:10.1038/nature08899 (2010).

30 Grimes, K. M. et al. The naked mole-rat exhibits an unusual cardiac myofilament protein profile providing new insights into heart function of this naturally subterranean rodent. Pflugers Arch 469, 1603–1613, doi:10.1007/s00424-017-2046-3 (2017).

31 Park, T. J. et al. Fructose-driven glycolysis supports anoxia resistance in the naked mole-rat. Science 356, 307–311, doi:10.1126/science.aab3896 (2017).

32 Nguyen, N. U. N. et al. A calcineurin-Hoxb13 axis regulates growth mode of mammalian cardiomyocytes. Nature 582, 271–276, doi:10.1038/s41586-020-2228-6 (2020).

33 Nguyen, P. D. et al. Interplay between calcium and sarcomeres directs cardiomyocyte maturation during regeneration. Science 380, 758–764, doi:10.1126/science.abo6718 (2023).

34 Mahmoud, A. I., Porrello, E. R., Kimura, W., Olson, E. N. & Sadek, H. A. Surgical models for cardiac regeneration in neonatal mice.

35 Aballo, T. J. et al. Ultrafast and Reproducible Proteomics from Small Amounts of Heart Tissue Enabled by Azo and timsTOF Pro. Journal of Proteome Research 20, 4203–4211, doi:10.1021/acs.jproteome.1c00446 (2021).

36 Meier, F. et al. diaPASEF: parallel accumulation–serial fragmentation combined with data-independent acquisition. Nature Methods 17, 1229–1236, doi:10.1038/s41592-020-00998-0 (2020).

37 Demichev, V. et al. dia-PASEF data analysis using FragPipe and DIA-NN for deep proteomics of low sample amounts. Nature Communications 13, 3944, doi:10.1038/s41467-022-31492-0 (2022).

38 Wieczorek, S. et al. DAPAR & ProStaR: software to perform statistical analyses in quantitative discovery proteomics. Bioinformatics 33, 135–136, doi:10.1093/bioinformatics/btw580 (2017).

39 Zhang, X. et al. Proteome-wide identification of ubiquitin interactions using UbIA-MS. Nature Protocols 13, 530–550, doi:10.1038/nprot.2017.147 (2018).

40 Ignatiadis, N., Klaus, B., Zaugg, J. B. & Huber, W. Data-driven hypothesis weighting increases detection power in genome-scale multiple testing. Nature Methods 13, 577–580, doi:10.1038/nmeth.3885 (2016).

41 Tucholski, T. et al. Distinct hypertrophic cardiomyopathy genotypes result in convergent sarcomeric proteoform profiles revealed by top-down proteomics. Proceedings of the National Academy of Sciences 117, 24691–24700, doi:doi:10.1073/pnas.2006764117 (2020).

42 Larson, E. J. et al. MASH Native: a unified solution for native top-down proteomics data processing. Bioinformatics 39, btad359, doi:10.1093/bioinformatics/btad359 (2023).

43 Kassambara, A. rstatix: Pipe-friendly framework for basic statistical tests. R package version 0.*6*. *0* (2020).

44 Hadley, W. ggplot2: Elegant Graphics for Data Analysis. (Springer-Verlag, 2016).

45 Kassambara, A. & Kassambara, M. A. Package ‘ggpubr’. R package version 0.*1* **6** (2020).

