## Supplementary Figures for "Integrated Proteomics Identifies Troponin I Isoform Switch as a Regulator of a Sarcomere-Metabolism Axis During Cardiac Regeneration"

### SUPPLEMENTAL MATERIAL: TABLE OF CONTENTS


***Section*** ***Pages***

Supplemental Figures....……………………………………………………...…....…………..3–10

Supplemental Tables....………………………………………………...…....…………..…...11

### Supplemental Figures

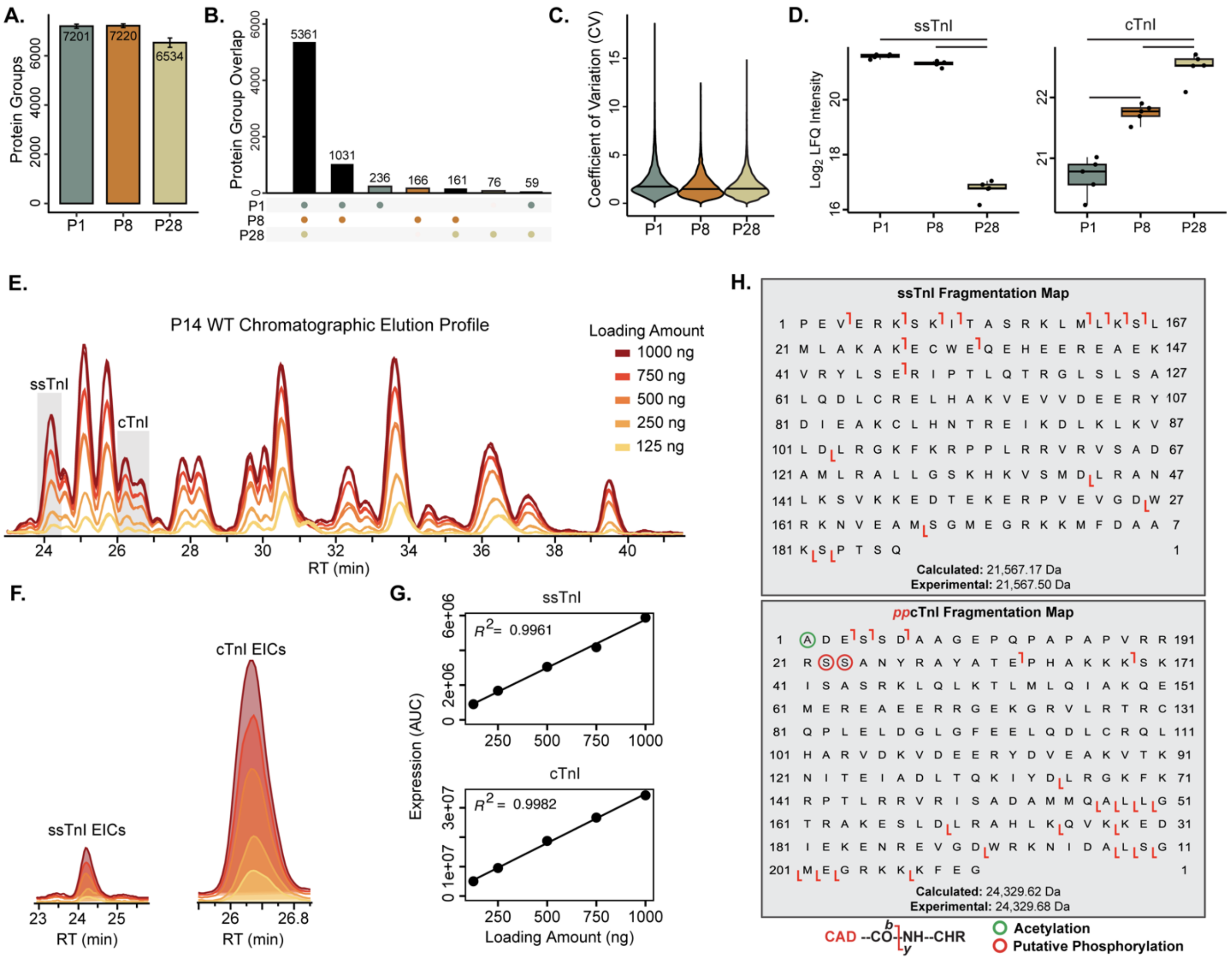


**Figure S1. Supplemental data from the proteomic analysis of postnatal mouse heart development. A.** Bar plot representing the average number of protein groups identified by global bottom-up proteomics from P1, P8, and P28 WT mouse hearts (n = 5 per group, error bars represent S.E.M.). **B.** UpSet plot of protein identifications between sample groups demonstrating conserved core proteome between sample groups. Protein group identifications were filtered to require n ≥ 3 per sample group for representation (n = 5 per group). **C.** Violin plot displaying extremely low coefficients of variations for protein groups quantifications. Line indicates median (n = 5 per group). **D.** Boxplots displaying ssTnI (Tnni1) and cTnI (Tnni3) expression throughout postnatal development (n = 5 per group, boxplots display median, upper, and lower quartile; dots represent individual hearts; bars between groups represent an adjusted p-value ≤ 0.05). **E.** Overlay of base peak chromatograms (BPCs) from three injection replicates of increasing amounts of total myofilament protein demonstrating reproducible chromatography and intensity between replicate injections. **F.** Representative extracted ion chromatograms (EICs) of ssTnI and cTnI generated from injection of 125, 250, 500, 750, and 1000 ng. EICs were generated from the 3-5 most abundant charge state ions from all proteoforms for each protein and averaged between three injection replicates. **G.** Quantification of the R2 value by plotting the area under the curve (AUC)

from EICs by the total protein loaded and performing a simple linear regression. **H.** Fragmentation maps for ssTnI (top) and ppcTnI (bottom) after collisionally activated dissocation (CAD) of MS1 precursor ions. Phosphorylation of serine 22 and 23 assigned based on fragmentation patterns and previously reported literature supporting phosphorylation of these residues.

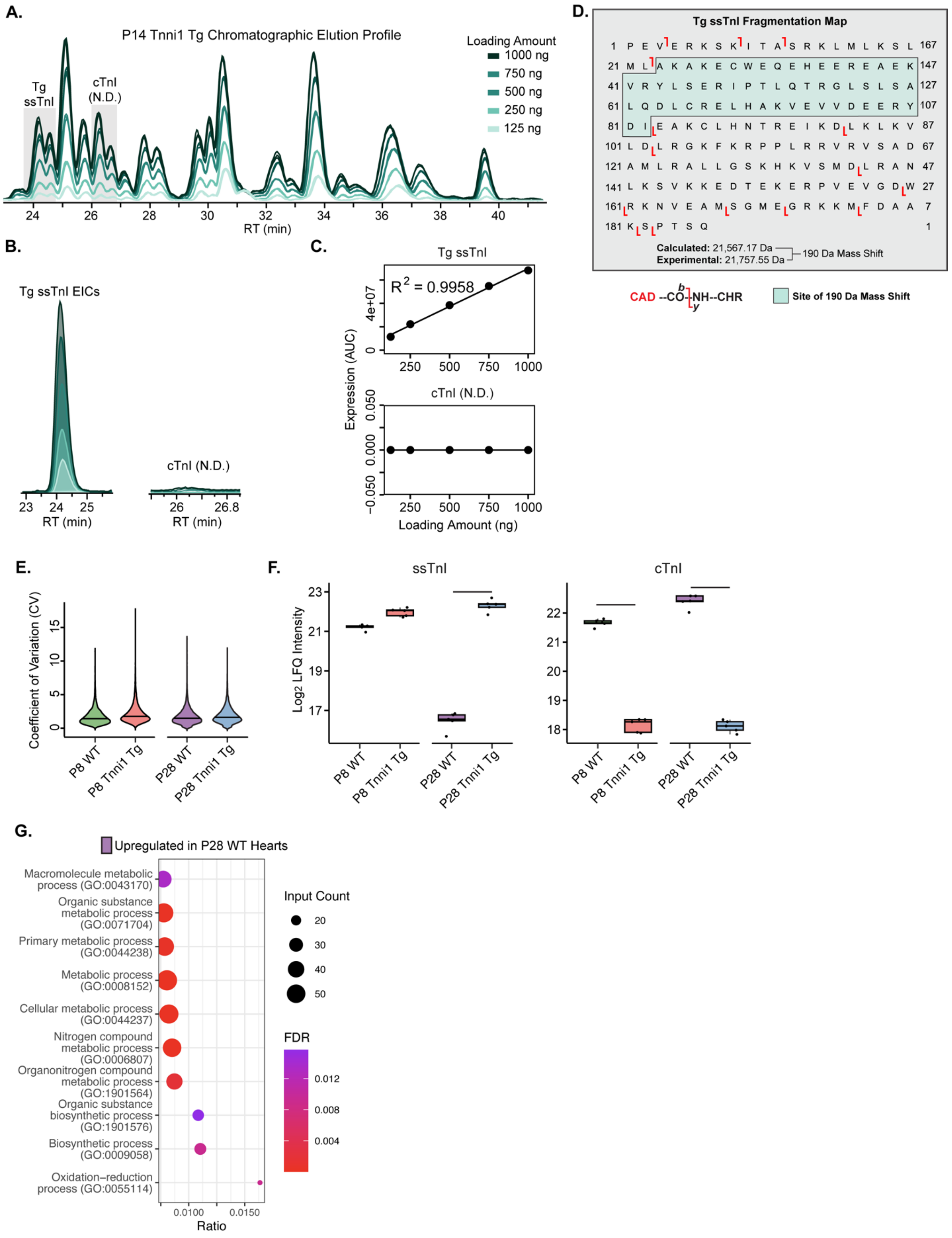
A.
P14 Tnni1 Tg Chromatographic Elution Profile
Loading Amount
1000 ng
750 ng
500 ng
250 ng
125 ng
Tg ssTnI
cTnI (N.D.)
RT (min)
B.
Tg ssTnI EICs
cTnI (N.D.)
RT (min)
C.
Tg ssTnI
R² = 0.9958
cTnI (N.D.)
Expression (AUC)
Loading Amount (ng)
D.
Tg ssTnI Fragmentation Map
Calculated: 21,567.17 Da
Experimental: 21,757.55 Da
190 Da Mass Shift
CAD --CO-NH--CHR
Site of 190 Da Mass Shift
E.
Coefficient of Variation (CV)
P8 WT
P8 Tnni1 Tg
P28 WT
P28 Tnni1 Tg
F.
ssTnI
cTnI
Log2 LFQ Intensity
G.
Upregulated in P28 WT Hearts
Macromolecule metabolic process (GO:0043170)
Organic substance metabolic process (GO:0071704)
Primary metabolic process (GO:0044238)
Metabolic process (GO:0008152)
Cellular metabolic process (GO:0044237)
Nitrogen compound metabolic process (GO:0006807)
Organonitrogen compound metabolic process (GO:1901564)
Organic substance biosynthetic process (GO:1901576)
Biosynthetic process (GO:0009058)
Oxidation−reduction process (GO:0055114)
Ratio
Input Count
FDR

**Figure S2. Supplemental data from the proteomic analysis of Tnni1 Tg and WT postnatal mouse hearts. A.** Overlay of BPCs from three injection replicates of increasing amounts of total myofilament protein demonstrating reproducible chromatography and intensity between replicate injections. **B.** Representative EICs of ssTnI and cTnI generated from injection of 125, 250, 500, 750, and 1000 ng. EICs were generated from the 3-5 most abundant charge state ions from all proteoforms for each protein and averaged between three injection replicates. cTnI was not detected (N.D.) in any of the injections **C.** Quantification of the R2 value by plotting the area under the curve (AUC) from EICs by the total protein loaded and performing a simple linear regression. **D.** Fragmentation maps for the Tnni1 Tg ssTnI after collisionally activated dissocation (CAD) of MS1 precursor ions. The Tnni1 Tg ssTnI displayed a +190 Da mass shift compared to WT ssTnI. As the unmodified ssTnI is not detected in any abundance, we attribute this mass shift to a mutation (likely on two residues) to the site highlighted in green. **E.** Violin plot displaying extremely low coefficients of variations for protein groups quantifications. Line indicates median ($n = 5$ per group). **F.** Boxplots displaying ssTnI (Tnni1) and cTnI (Tnni3) expression in WT and Tnni1 Tg mouse hearts at P8 and P28 ($n = 5$ per group, boxplots display median, upper, and lower quartile; dots represent individual hearts; bars between groups represent an adjusted p-value $\leq 0.05$). **G.** Selected STRING biological process gene ontology (GO) plots of the proteins upregulated in P28 WT hearts compared to P28 Tnni1 Tg hearts. There were not enough differentially expressed proteins upregulated in P28 Tnni1 Tg hearts to perform GO analysis. Ratio represents the fractions of all proteins in the GO category that were identified. Dot size corresponds to the number of identified proteins within a GO category. Color represents FDR-adjusted p-value of the overrepresentation test.

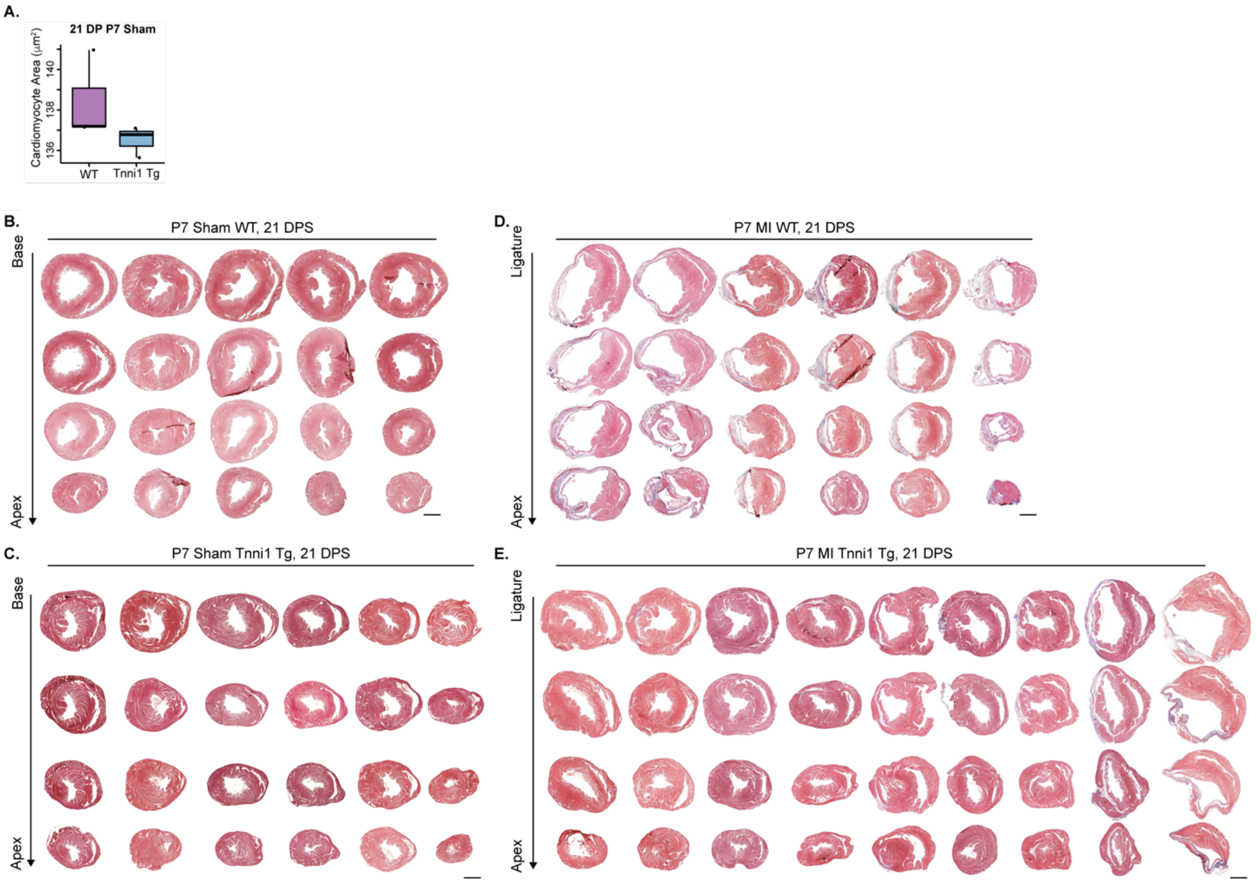


**Figure S3. Supplemental data from the analysis of injury response in P7 WT and Tnni1 Tg mice. A.** P28 hearts (21 DP P7 sham surgery) display similarly sized cardiomyocytes (n = 3 per group, boxplots display median, upper, and lower quartile; dots represent individual hearts; no significant difference detected as determined by a 2-tailed unpaired Student's *t*-test). **B-E.** Masson's trichrome-stained heart sections of WT **(B)** and Tnni1 Tg **(C)** mice hearts 21 DP P7 sham surgery and sections of WT **(D)** and Tnni1 Tg **(E)** mice hearts 21 DP P7 MI surgery. Serial sections were cut from the base to the apex of the heart. All hearts are shown. Scale bars, 1 mm.

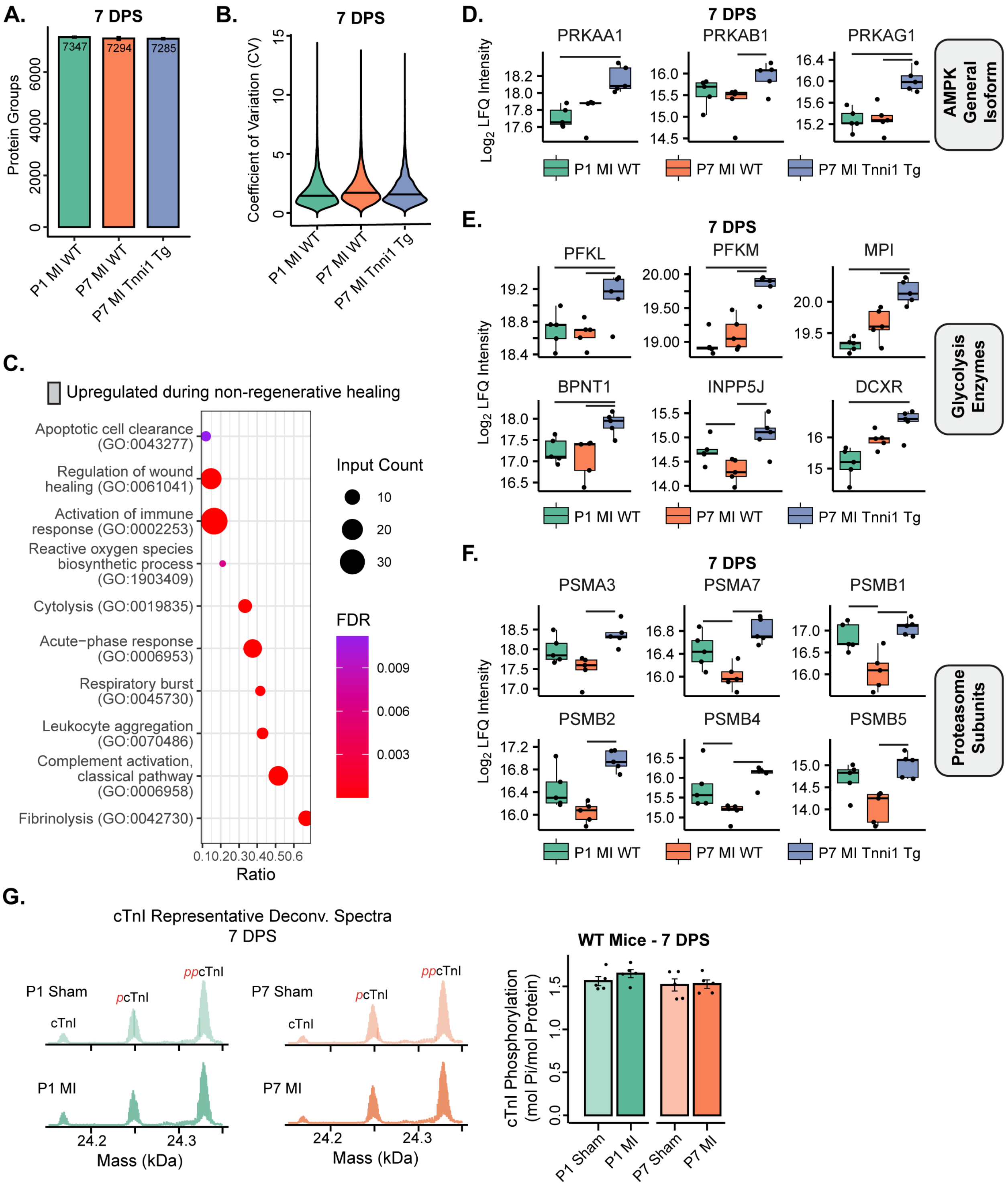


**Figure S4. Supplemental data examining the proteomic profiles of WT and Tnni1 Tg mice 7 days-post MI. A.** Bar plot representing the average number of protein groups identified by global bottom-up proteomics from P1 MI WT, P7 MI WT, and P7 MI Tnni1 Tg mouse hearts 7 DPS (n = 5 per group, error bars represent S.E.M.). **B.** Violin plot displaying extremely low coefficients of variations for protein groups quantifications. Line indicates median (n = 5 per group). **C.** Selected

STRING biological process gene ontology (GO) plots of the proteins that are elevated during WT non-regenerative. Ratio represents the fractions of all proteins in the GO category that were identified. Dot size corresponds to the number of identified proteins within a GO category. Color represents FDR-adjusted p-value of the overrepresentation test. **D-F.** Boxplots of selected proteins upregulated during Tnni1 Tg regenerative healing, including every subunit of the general isoform of AMPK **(D)**, additional proteins related to glycolysis and glycogen utilizing **(E)**, and additional subunits in the proteasome **(F)** (n = 5 per group, boxplots display median, upper, and lower quartile; dots represent individual hearts; bars between groups represent an adjusted p-value ≤ 0.05). **G.** Top-down proteomic analysis of cTnI phosphorylation 7 DPS during regenerative and non-regenerative responses to injury. Left: Representative deconvoluted spectra of cTnI 7 DPS. Right: Quantification of relative expression of phosphorylated cTnI to total cTnI expression. Bars represent average, error bars represent standard error of the mean (S.E.M.), dots represent individual data points. No statistical difference between the groups as determined by a 2-tailed unpaired Student's *t*-test.

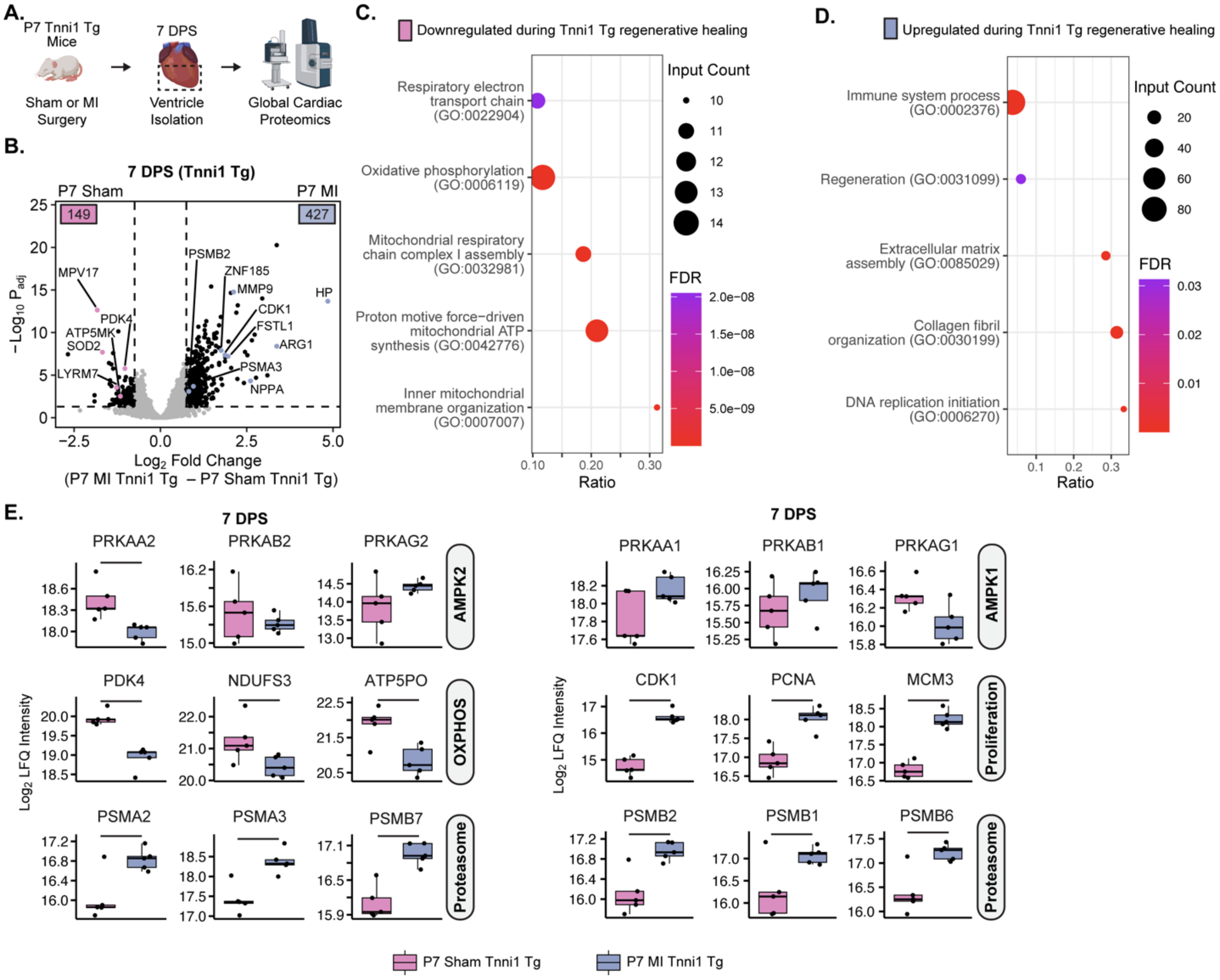


**Figure S5. Proteomic analysis of injury response compared to sham hearts in Tnni1 Tg mice.** **A.** Workflow of experimental design. Sham and myocardial infarction surgeries were performed on and P7 Tnni1 Tg mice (n = 5 per group). 7 days post-surgery (DPS), hearts were harvested for global bottom-up cardiac proteomics. **B.** Volcano plot demonstrating fold-change in protein expression between sham and MI Tnni1 Tg hearts 7 DPS in P7 mice. The number of significantly upregulated proteins per group is shown in the bottom corners of each comparison (n = 5 per group). Metabolic, mitotic, and proteasomal proteins are highlighted (adjusted p-value ≤ 0.05 and |Log2 Fold Change| ≥ 0.75 significance thresholds). **C and D.** Selected STRING biological process gene ontology (GO) plots of the proteins that are downregulated **(C)** and upregulated **(D)** during Tnni1 Tg regenerative healing compared to sham hearts. Ratio represents the fractions of all proteins in the GO category that were identified. Dot size corresponds to the number of identified proteins within a GO category. Color represents FDR-adjusted p-value of the overrepresentation test. **E.** Protein expression boxplots of selected proteins related to AMPK, oxidative phosphorylation (OXPHOS), proliferation, and the proteasome during the Tnni1 Tg regenerative response to injury compared to sham animals (n = 5 per group, boxplots display median, upper, and lower quartile; dots represent individual hearts; bars between groups represent an adjusted p-value ≤ 0.05).

**Supplemental Tables**

**Table S1. General experimental information and database search results.** A supplemental table with sheets detailing experimental design for proteomic experiments, protein group search results from diaNN, and peptide search results diaNN.

**Table S2. Supplemental information for investigation of postnatal cardiac development in mice.** A supplemental table with sheets detailing differentially expressed proteins between P1, P8 and P28 hearts, STRING gene ontology searches, ions used for EIC generation, MS/MS fragments for top-down protein identification, and statistical results for the top-down analyses.

**Table S3. Supplemental information for investigation of baseline proteome composition in P8 and P28 WT and Tnni1 Tg mice.** A supplemental table with sheets detailing differentially expressed proteins between P8 and P28 WT and Tnni1 Tg mice, STRING gene ontology searches, ions used for EIC generation, MS/MS fragments for top-down protein identification, and statistical results for the top-down analyses.

**Table S4. Statistical information for WT and Tnni1 Tg P7 MI injury response.** A supplemental table detailing the statistical results for the injury response analyses.

**Table S5. Supplemental information for investigation of proteomic analysis 7 days-post MI or sham surgery in Tnni1 Tg mice.** A supplemental table with sheets detailing differentially expressed proteins between sham and MI Tnni1 Tg mice, STRING gene ontology searches, ions used for EIC generation, and statistical results for the top-down analyses.
